# Polygenic Scores Predict the Development of Alcohol and Nicotine Use Problems from Adolescence through Young Adulthood

**DOI:** 10.1101/2020.07.29.227439

**Authors:** Joseph D. Deak, D. Angus Clark, Mengzhen Liu, C. Emily Durbin, William G. Iacono, Matt McGue, Scott I. Vrieze, Brian M. Hicks

## Abstract

**Objective:** Molecular genetic studies of alcohol and nicotine have identified many genome-wide loci. We examined the predictive utility of drinking and smoking polygenic scores (PGS) for alcohol and nicotine use from late childhood to early adulthood, substance-specific versus broader-liability PGS effects, and if PGS performance varied between consumption versus pathological use.

**Methods:** Latent growth curve models with structured residuals were used to assess the predictive utility of drinks per week and regular smoking PGS for measures of alcohol and nicotine consumption and problematic use from age 14 to 34. PGSs were generated from the largest discovery sample for alcohol and nicotine use to date (i.e., GSCAN), and examined for associations with alcohol and nicotine use in the Minnesota Twin Family Study (N=3225).

**Results:** The drinking PGS was a significant predictor of age 14 problematic alcohol use and increases in problematic use during young adulthood. The smoking PGS was a significant predictor for all nicotine use outcomes. After adjusting for the effects of both PGSs, the smoking PGS demonstrated incremental predictive utility for most alcohol use outcomes and remained a significant predictor of nicotine use trajectories.

**Conclusions:** Higher PGS for drinking and smoking were associated with more problematic levels of substance use longitudinally. The smoking PGS seems to capture both nicotine-specific and non-specific genetic liability for substance use, and may index genetic risk for broader externalizing behavior. Validation of PGS within longitudinal designs may have important clinical implications should future studies support the clinical utility of PGS for substance use disorders.

## Introduction

Alcohol and nicotine use, respectively, contribute to 3 million (5.3%) and 7 million (12.3%) deaths worldwide each year, making both leading causes of global mortality [1,2]. While public health interventions have reduced some negative consequences associated with alcohol and nicotine use, a significant portion of the population continue to meet criteria for nicotine use disorder (NUD) and alcohol use disorder (AUD), suggesting that genetic influences likely contribute to problem use for important subgroups. Twin studies report heritability estimates of approximately 50% for AUD [3] and NUD [e.g.,4], and large consortia of genome-wide association studies (GWAS) have identified hundreds of loci that exhibit genome-wide significant associations with alcohol and nicotine use phenotypes [5–7], providing new avenues for research on the genetic influences on substance use.

Polygenic scores (PGS) are one method for modeling aggregate genetic risk across the genome, and have provided valuable information about the unique and shared genetic influences on alcohol and nicotine use. PGS can be generated from a GWAS discovery sample by weighting SNPs relative to the strength of their association with a given phenotype to calculate a measure of individual genetic risk in a target sample. For example, PGS calculated from GWAS-identified associations for alcohol use have predicted alcohol-related outcomes in independent samples [8,9]. PGS for alcohol and nicotine use have also predicted use of a variety of other substances (e.g., cannabis, cocaine, amphetamines, ecstasy, hallucinogens [5,10]), suggesting these PGS index non-specific genetic influences on substance use.

While studies using PGS are beginning to trace the contours of the genetic architecture of substance use, they have yet to examine the influence of aggregate genetic risk on patterns of substance use over time. This is an important next step, because alcohol and nicotine use exhibit strong age-related mean-level trends, with typical initiation in adolescence followed by peak use in young adulthood and normative declines in heavy use and substance use disorders by age 30 [11]. Understanding the etiology of substance use then requires accounting for these normative patterns of emergence, escalation, and decline, and there is some evidence that genetic influences for substance use varies across development [12,13].

Initial efforts using PGSs to predict alcohol and nicotine use trajectories over time have had some success. A PGS for cigarettes smoked per day predicted later cigarette smoking and NUD in early adulthood [14,15], but not alcohol use [14], suggesting the PGS measured substance-specific genetic influences on nicotine use. Evidence for the predictive utility of alcohol-related PGS has been mixed. One study found that a PGS for AUD was associated with levels of alcohol use in males at age 15.5 and greater increases of alcohol use at age 21.5 [16], while other studies predicting alcohol use in college student drinkers over time have returned both positive [17] and null results [18]. Most prior studies were limited by smaller GWAS discovery samples relative to the much larger recent GWAS consortia of alcohol and nicotine use. Additionally, the longitudinal studies of alcohol use-related PGS primarily examined college student populations assessed across a four-year timespan during which environmental influences are enriched for substance use, potentially limiting the influence of polygenic contributions in this context.

We address these limitations using PGS measures derived from the GWAS & Sequencing Consortium of Alcohol and Nicotine use (GSCAN), the largest GWAS discovery sample for alcohol and nicotine use to date [5], and examined their utility to predict trajectories of alcohol and nicotine use and problem use from late childhood through young adulthood (i.e., age 14 – 34). Strengths of this approach include the ability to make stronger inferences about when in the developmental progression of substance use (e.g., initiation of use, escalation of use) these genetic influences have their effects, and the long follow-up period ensures that polygenic influences for alcohol and nicotine use are likely to have been expressed for most people. The present study also aimed to examine whether the predictive utility of respective alcohol and nicotine-related PGS were limited to their specific substance or generalized to predict trajectories of both alcohol and nicotine use. Given prior evidence suggesting differences in the genetic architecture of alcohol use versus AUD [19,20], a final aim was to examine whether the predictive utility of GSCAN alcohol and smoking PGSs differed for consumption outcomes in comparison to symptoms of AUD and NUD.

## Methods

### Participants

Participants were members of the Minnesota Twin Family Study (MTFS), a longitudinal study of 3762 (52% female) twins (1881 pairs) investigating the development of substance use disorders and related conditions [21–23]. All twin pairs were the same sex and living with at least one biological parent within driving distance to the University of Minnesota laboratories at the time of recruitment. Exclusion criteria included any cognitive or physical disability that would interfere with study participation. Twins were recruited the year they turned either 11-years old (*n*=2510; the younger cohort) or 17-years old (*n*=1252; the older cohort). Twins in the younger cohort were born from 1977 to 1984 and 1988 to 1994, while twins in the older cohort were born between 1972 and 1979. Families were representative of the area they were drawn from in terms of socioeconomic status, mental health treatment history, and urban vs rural residence [21]. Consistent with the demographics of Minnesota for the target birth years, 96% of participants reported non-Hispanic White race and ethnicity.

The younger cohort was assessed at ages 11 (M_age_=11.78 years; SD=0.43 years) and 14 (M_age_=14.90 years; SD=0.31 years), and all twins were assessed at target ages 17 (M_age_=17.85 years; SD=0.64 years), 21 (M_age_=21.08 years; SD=0.79 years), 24 (M_age_=24.87 years; SD=0.94 years), and 29 (M_age_=29.43 years; SD=0.67 years). A subgroup of twins from the younger cohort were also assessed at age 34 (M_age_=34.62 years; SD=1.30 years). Supplemental Table 1 provides the number of participants for each assessment and descriptive statistics for the study measures. Participation rates ranged from 80% to 93% among those recruited for a given assessment. The total sample included 1205 monozygotic (51.5% female) and 676 dizygotic (52.8% female) twin pairs [21,24].

#### Alcohol Use and AUD

All alcohol and nicotine variables were assessed during structured clinical interviews, while the use variables were also assessed using a computerized self-report questionnaire at ages 11, 14, and 17 that was completed in private. Alcohol variables included the average number of drinks per occasion in the past 12 months (i.e., alcohol quantity), *DSM-III-R* symptoms of alcohol abuse and dependence (the diagnostic system when the study began, hereafter referred to as AUD symptoms), and an alcohol problems composite variable calculated at each age consisting of the average of mean alcohol quantity, AUD symptoms, and maximum number of drinks consumed in 24 hours (i.e., max drinks). Free responses to alcohol quantity and max drinks, as well as the number of alcohol abuse and dependence symptoms were converted to scales that ranged from 0 to 8. In terms of problem use, the lifetime prevalence of *DSM-III-R* AUD (3 or more symptoms of abuse or dependence) was 26%.

#### Nicotine Use and NUD

Nicotine variables included average quantity per day (e.g., cigarettes smoked per day), *DSM-III-R* symptoms of nicotine dependence (hereafter referred to as NUD symptoms), and an age-specific nicotine use problems composite variable calculated from the averages of mean nicotine quantity, NUD symptoms, and typical frequency of nicotine use (i.e., number of days per month) in the past 12 months. Free responses were converted to a 0 to 4 scale for nicotine frequency and a 0 to 6 scale for nicotine quantity and NUD symptoms. The lifetime prevalence of *DSM-III-R* NUD was 33%.

#### PGS Methods

PGS were generated from the GSCAN discovery sample using GWAS summary statistics for drinks per week and ever being a regular smoker, following removal of the MTFS sample to avoid overlap with the target sample [5]. PGS were created for participants of European ancestry, confirmed via principal components analysis [25], in the MTFS target sample following imputation to the most recent Haplotype Reference Consortium reference panel [26] and restricted to autosomal HapMap3 variants with a minor allele frequency (MAF) ≥ 0.01 and an imputation quality > 0.7. The resulting filtered variants (i.e., ~1 million variants) were then submitted to LDpred [27], given the familial structure of the target sample, to generate beta weights in the MTFS sample, including variants of all significance levels (i.e., *p*-value threshold ≤ 1). Individual PGS were then calculated in PLINK 1.9 [25] for all individuals meeting inclusion criteria for the present study (*N*=3225). Given the study aim of examining substance-specific and cross-substance influences of both alcohol and nicotine use PGS, GSCAN drinks per week (N_full-sample_ = 941,280) and smoking initiation (N_full-sample_ = 1,232,091) were selected as PGS variables as they are the largest, and most comparably powered, variables available in the GSCAN discovery sample; and thus, aptly suited to examine the primary questions of interest in the present study.

### Data Analytic Strategy

Latent growth models with structured residuals (LGM-SR; see Figure 1) were used to model developmental trends in the alcohol and nicotine use outcomes [28,29]. These models include intercept factors that reflect status at the first time point (age 14 as there was almost no substance use at age 11), and slope factors that reflect the rate of change over the course of the study. Slope factors were specified using a latent basis approach. That is, the first and last basis coefficient were fixed to 0 and 1, respectively, and the intervening coefficients were estimated, which provides a parsimonious way of capturing non-linear trajectories [30]^1^. Intercept and slope factors were allowed to vary to capture individual differences in growth. The residual structure included occasion-specific latent factors that account for deviations from the intercept and slope implied trajectories. The autoregressive paths linking adjacent residual factors capture associations between variables over time after accounting for general growth trend (**Figure 1**) and were included because not accounting for residual autoregressive effects can lead to biased variance estimates in the growth factors [31,32].

**Figure 1.**
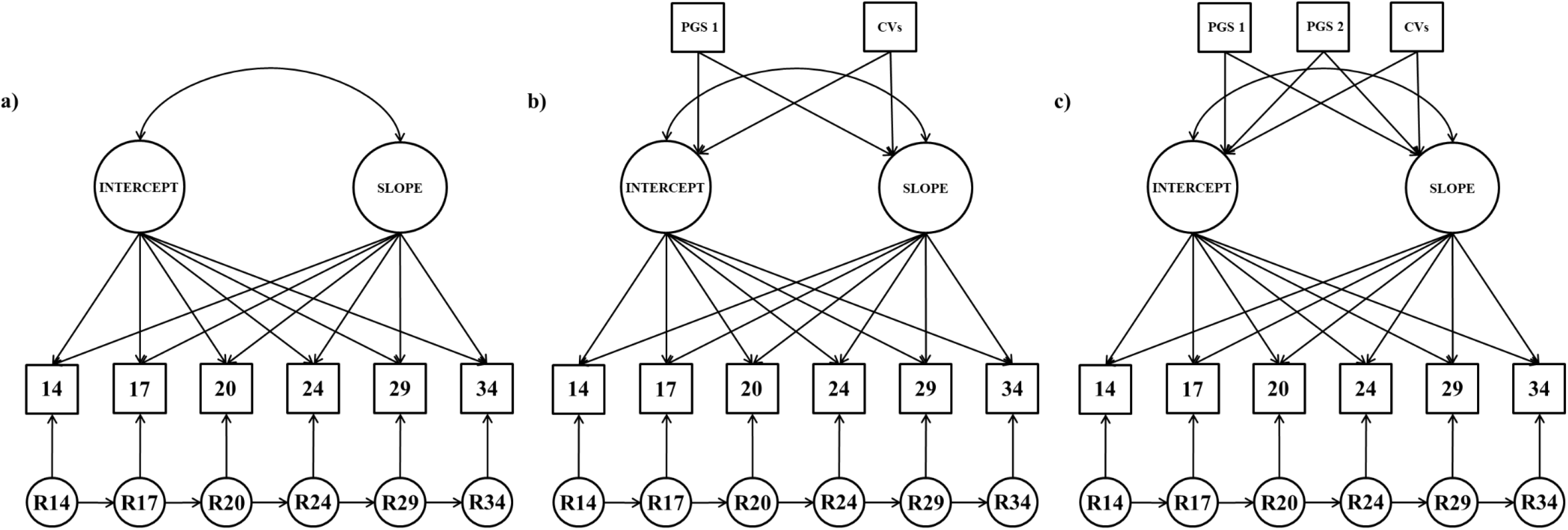
Unconditional and Conditional Latent Growth Models with Structured Residuals. Panel a depicts the unconditional latent growth model; Panel b depicts the single PGS conditional latent growth model; Panel c depicts the two PGS conditional latent growth model. R=residual factor; PGS=polygenetic risk score; CVs=covariates (first 5 eigenvalues and sex). Variances and mean structure omitted from figure for clarity of presentation.

Unconditional LGM-SR models were first fit to each outcome (Figure 1-Panel a). Conditional models were then fit in which the growth factors were regressed on a single PGS and the control variables (Single PGS Predictor Model; Figure 1-Panel b). The control variables included participant sex and the first five genetic principal components [33] to adjust for underlying ancestral substructure. Finally, conditional models were estimated in which the growth factors were regressed on both PGS’ simultaneously along with the control variables (Two PGS Predictor Model; Figure 1-Panel c). All major analyses were conducted using Mplus v8.4 [34] with full information maximum likelihood estimation [35]. Confidence intervals were derived using clustered (by family) percentile bootstrapping (with 1000 draws), which performs well when estimating confidence intervals with skewed variables such as substance use, and accounts for the non-independence of the observations (i.e., twins nested within families) [36].

## Results

Descriptive information for the study variables is reported in the supplemental material (OSF). Mean-levels of the alcohol and nicotine use outcomes increased from age 11 to age 20, and then decreased from age 20 to age 34. The rank-order stability of the alcohol and nicotine use outcomes between adjacent time points ranged from *r*=.33 to .83 (mean *r*=.57). Models were fit both with and without participants that consistently abstained from substance use across time. Conclusions were similar across these models, and so we report the results for models fit using the full sample.

The univariate models for alcohol and nicotine use related outcomes all fit the data well by conventional standards [37]. Parameter estimates were consistent with the observed trajectories, suggesting a rise in alcohol and nicotine use throughout adolescence, and then a gradual decline in values after age 20. There was a statistically significant degree of variability in all of the growth factors (see OSF).

### Alcohol Use Outcomes

Standardized path coefficients from the single PGS conditional models can be found in **Table 1**. The drinks per week PGS was a statistically significant predictor of small magnitude (mean β=.09) for all alcohol use-related growth factors except for the alcohol quantity intercept, though the effect size was similar (β=.10). The smoking PGS was a statistically significant predictor of small to medium magnitude (mean β=.17) for all alcohol use-related growth factors except for the alcohol quantity slope (β=.05).

**Table 1.** Standardized Coefficients to Intercept and Slope Factors From Single and Double PGS Predictor Models

|  | Drinks Per Week PGS |  |  |  | Regular Smoking PGS |  |  |  |
| --- | --- | --- | --- | --- | --- | --- | --- | --- |
|  | Intercept<br>(1 PGS) | Intercept<br>(2 PGS) | Slope<br>(1 PGS) | Slope<br>(2 PGS) | Intercept<br>(1 PGS) | Intercept<br>(2 PGS) | Slope<br>(1 PGS) | Slope<br>(2 PGS) |
| Alcohol Use Disorder | **.12**<br>[.06, .22] | **.10**<br>[.04, .18] | **.06**<br>[.01, .11] | .04<br>[-.02, .09] | **.12**<br>[.06, .25] | **.10**<br>[.04, .22] | **.12**<br>[.07, .17] | **.11**<br>[.06, .16] |
| Alcohol Quantity | .10<br>[-.04, .48] | .01<br>[-.17, .27] | **.07**<br>[.01, .15] | .07<br>[.00, .14] | **.36**<br>[.19, 1.32] | **.35**<br>[.19, 1.33] | .05<br>[-.02, .12] | .03<br>[-.04, .10] |
| Alcohol Composite | **.10**<br>[.01, .26] | .04<br>[-.05, .17] | **.08**<br>[.03, .13] | **.06**<br>[.01, .11] | **.25**<br>[.16, .53] | **.24**<br>[.15, .53] | **.11**<br>[.05, .16] | **.09**<br>[.04, .15] |
| Nicotine Use Disorder | **.10**<br>[.03, .18] | .06<br>[-.02, .13] | .04<br>[-.02, .10] | .00<br>[-.06, .06] | **.19**<br>[.12, .28] | **.17**<br>[.11, .27] | **.18**<br>[.12, .23] | **.18**<br>[.12, .24] |
| Nicotine Quantity | **.12**<br>[.03, .27] | .05<br>[-.05, .18] | .02<br>[-.04, .08] | .00<br>[-.07, .07] | **.30**<br>[.19, .57] | **.29**<br>[.18, .55] | **.09**<br>[.02, .16] | **.09**<br>[.02, .16] |
| Nicotine Composite | **.11**<br>[.03, .22] | .06<br>[-.03, .15] | .03<br>[-.02, .09] | .00<br>[-.06, .05] | **.23**<br>[.15, .38] | **.22**<br>[.14, .35] | **.18**<br>[.12, .24] | **.18**<br>[.12, .24] |

1 PGS=coefficients from single PGS predictor model; 2 PGS=coefficients from double PGS predictor model; Bold=95% confidence interval does not include 0. Only one PGS was entered as a predictor in each single PGS predictor model; both PGSs were entered as predictors simultaneously in the double PGS predictor model. Eigenvalues 1 through 5 and sex were entered into each model along with the PGS as control variables; coefficients for control variables not presented. Confidence intervals derived via clustered non-parametric percentile bootstrap with 1,000 draws.

Standardized path coefficients from the two PGS conditional models can be found in Table 1. Most of the effects for the drinks per week PGS were not significant and negligible in magnitude (mean β=.05), except for the statistically significant but small effects on the AUD intercept factor (β=.10) and alcohol composite slope factor (β=.06). The smoking PGS remained a statistically significant predictor of small to medium effect for all the alcohol use-related growth factors (mean β=.15) except for the alcohol quantity slope (β=.03).

### Nicotine Use Outcomes

The smoking PGS was a statistically significant predictor of small to medium magnitude (mean β=.21) for all nicotine use-related growth factors. The drinks per week PGS was a modest, statistically significant predictor (mean β=.11) of all of the intercept factors, but none of the slope factors (mean β=.03). In the two PGS conditional models, the smoking PGS remained a statistically significant predictor of small to medium effect for all nicotine use-related growth factors (mean β=.18). None of the drinks per week PGS effects remained significant (mean β=.03). Substance use growth trajectories for individuals with high (i.e., 1.5 standard deviation above the mean), low (i.e., 1.5 standard below the mean), and average scores for the respective PGSs can be found in Figure 2.

**Figure 2.**
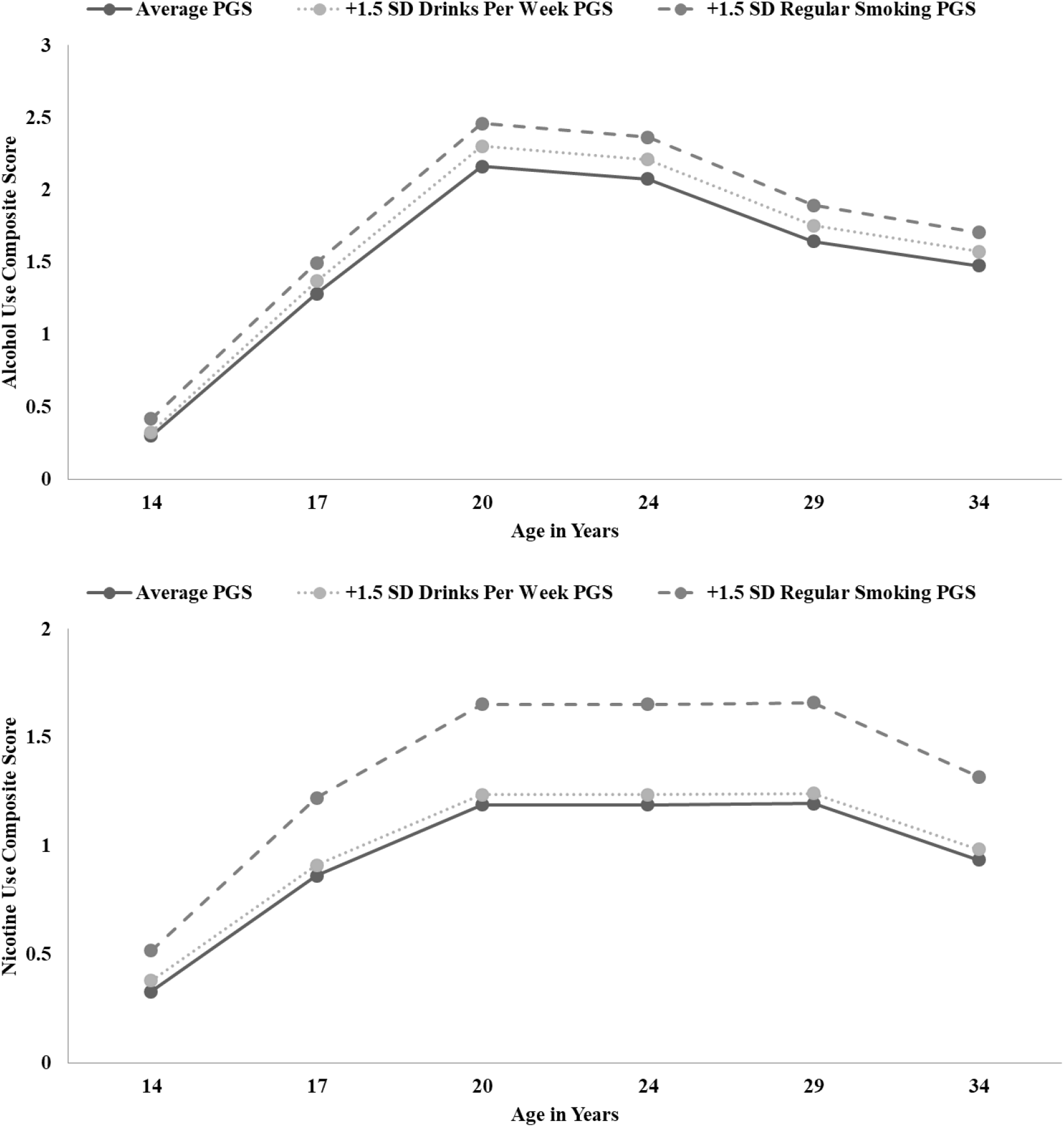
Growth Trajectories from the Two PGS Predictor Models for Alcohol and Nicotine Use Composites. Growth trajectories for the alcohol composite presented in the top panel, growth trajectories for the nicotine composite presented in the lower panel. Age in years presented on X axis, composite scores presented on Y axis. Trajectories are based on the parameter estimates from the full two PGS predictor models. The three lines depict trajectories for those with average scores on both PGSs (solid line), high scores (1.5 standard deviation above the mean) on the drinks per week PGS and average scores on the regular smoking PGS (dotted line), and average scores on the drinks per week PGS and high scores on the regular smoking PGS (dashed line).

We also fit models that regressed scores for each growth factor (e.g., AUD slope) on the same predictors of the two PGS conditional models, but also added scores for the comparable growth factor of the other substance (e.g., NUD slope). Results were consistent with those of the two PGS conditional models (see online supplement for details) indicating the PGS retained predictive utility even after adjusting for the phenotypic overlap between alcohol and nicotine use.

## Discussion

We extended prior studies investigating genetic influences for alcohol and nicotine use by examining the utility of PGSs to predict the developmental progression of alcohol and nicotine use. Using a longitudinal design, we found that higher drinks per week and smoking PGSs were each associated with more problematic levels of use of the respective substances in middle adolescence, and a greater rate of increase in alcohol and nicotine use problems through early adulthood. These findings demonstrate that alcohol and nicotine use-related PGS are robust predictors of trajectories for problematic alcohol and nicotine use, and suggest the potential for clinical applications utilizing polygenic risk profiles to help inform intervention strategies.

We also examined differences in prediction for regular use versus symptoms of substance use disorder. Though most effects were consistent across regular use and substance use disorder symptoms, we found that the drinks per week and smoking PGSs were more predictive of AUD symptoms relative to average quantity of alcohol consumed per drinking occasion, though each PGS was predictive of all substance-specific measures over time. Prior studies have reported substantial, but incomplete genetic overlap between alcohol consumption and AUD (*r_g_ ~*0.7) [38], and these shared genetic influences do not seem to extend to other psychiatric conditions. Notably, AUD tends to have moderate and robust genetic correlations with other psychiatric phenotypes, while measures of alcohol consumption have smaller and often non-significant genetic correlations with psychiatric disorders [7,19]. Future studies are needed to better decipher the common and unique genetic influences on normative versus problematic use, and how this overlap may vary over time.

Additionally, we examined the substance-specific versus generalized effects of the PGS, as well as their incremental predictive utility. We found that the smoking PGS was a stronger predictor of both nicotine and alcohol use-related phenotypes, even after adjusting for its overlap with the drinks per week PGS. In contrast, the predictive utility of the drinks per week PGS was mostly specific to alcohol use, and even most of these associations were non-significant after adjusting for its overlap with the regular smoking PGS. Thus, the smoking PGS seems to measure both nicotine-specific and non-specific genetic liability for substance use problems, potentially serving as an index of externalizing behavior more broadly (i.e., poor impulse control and norm violating behavior), while the drinks per week PGS seems to measure genetic risk that is relatively specific to alcohol.

The notion that smoking polygenic risk is broadly predictive of externalizing behaviors is supported by results from the initial wave of GSCAN showing that the smoking PGS was associated with a variety of substances (e.g., alcohol use, cannabis use, cocaine use [5]). The evidence for common genetic influences across substances is also consistent with prior multivariate twin studies that posited a common genetic etiology for engaging in externalizing behaviors [39–41]. Given these prior findings, the smoking PGS may index non-specific genetic risk for externalizing behaviors, which would account for its robust prediction of nicotine and alcohol use outcomes and its incremental predictive utility relative to the drinks per week PGS.

While an exciting potential, the clinical utility of polygenic scoring approaches for substance use and psychiatric traits requires substantial advancement before the utility of incorporating genetic profiles into treatment planning and prevention approaches can be realized. While stratifying individuals based upon “polygenic risk” has been shown to aid in mitigating adverse health outcomes related to some health conditions (e.g., coronary disease; 42), similar success has yet to be demonstrated for substance use disorders (SUD). Concerns related to the clinical utility of PGS for substance use outcomes include that current SUD PGS account for a relatively small proportion of variance for clinical phenotypes (~10% at best), misinterpretation of what a PGS means (95% percentile of genetic risk ≠ 95% likelihood of developing disorder), and discrimination of patients based upon genetic information [e.g., 43–45]. Given this context, findings from the current study provide initial evidence that polygenic influences are associated with increased rates of substance use and related problems over the span of important developmental periods (**Figure 2)**, periods in which targeted prevention and intervention strategies (e.g., psychoeducation, lifestyle changes) may help circumvent adverse substance-related outcomes later in life. As the field continues to evaluate the potential benefits, and drawbacks, of the clinical utility of PGS for substance use outcomes, considerations on how these approaches may be examined within longitudinal designs should be considered.

Limitations of the study include that the PGS were generated from large GWAS that were conducted in countries in which the population is primarily of European descent, and due to varying allele frequencies across ancestral groups, the degree to which the current results generalize to other ancestral groups is uncertain [46]. Notably, this limitation has the potential to proliferate health disparities with precision medicine efforts if these findings are only applicable to individuals of European ancestry, further prioritizing the importance of extending these efforts to diverse ancestry groups [47]. Additionally, genetic influences on substance use behaviors are influenced by a variety of environmental factors (e.g., peer influences), and genetic and environmental influences vary across development stages [14,48]. Thus, future studies examining how these PGS interact with environmental influences longitudinally are needed.

Despite these limitations, the current study represents a successful extension of prior work by validating the predictive utility of the PGS approach in longitudinal models of alcohol and nicotine use phenotypes across late childhood and early adulthood. The results also provide initial evidence that the regular smoking PGS may index non-specific genetic risk for substance use and externalizing behaviors in general. This validation serves as a key step in demonstrating polygenic prediction of alcohol and nicotine use outcomes across important developmental periods, and may serve as foundational should the incorporation of polygenic scoring approaches be supported by future studies examining the relative pros and cons of incorporating genetic information for prevention and intervention approaches aimed at circumventing negative substance use outcomes.

## Acknowledgement

This work was supported by United States Public Health Service grants R37 AA09367 (McGue), R01 AA024433 (Hicks), and T32 AA007477 (Blow) from the National Institute of Alcohol Abuse and Alcoholism and R01 DA034606 (Hicks), R37 DA005147 (Iacono), R01 DA013240 (Iacono), R01 DA044283 (Vrieze), R01 DA037904 (Vrieze), R01 DA042755 (McGue/Vrieze) and U01 DA046413 (Vrieze) from the National Institute on Drug Abuse.

## Footnotes

1 Alternative specifications of the growth model (e.g., piecewise models) were considered, and lead to the same conclusions as reported here

## References

1. World Health Organization, 2018. Global Status Report on Alcohol and Health 2018 Ed, World Health Organization, Geneva, Switzerland, 2018.

2. World Health Organization. WHO Report on the Global Tobacco Epidemic, 2017: Monitoring Tobacco Use and Prevention Policies. World Health Organization; 2017.

3. Verhulst B, Neale MC, Kendler KS. The heritability of alcohol use disorders: a meta-analysis of twin and adoption studies. Psychol Med. 2015 Apr;45(5):1061–72.

4. Agrawal A, Verweij KJH, Gillespie NA, et al. The genetics of addiction—a translational perspective. Translational psychiatry. 2012;2(7):e140–e140.

5. Liu M, Jiang Y, Wedow R, et al. Association studies of up to 1.2 million individuals yield new insights into the genetic etiology of tobacco and alcohol use. Nature genetics. 2019;51(2):237–244.

6. Johnson EC, Chang Y, Agrawal A. An update on the role of common genetic variation underlying substance use disorders. Curr Genetic Med. 2020; Rep 8:35–46

7. Sanchez-Roige S, Palmer AA, Clarke T-K. Recent efforts to dissect the genetic basis of alcohol use and abuse. Biological Psychiatry. 2020;87(7):609–618.

8. Salvatore JE, Aliev F, Edwards AC, et al. Polygenic scores predict alcohol problems in an independent sample and show moderation by the environment. Genes. 2014;5(2):330–346.

9. Savage JE, Salvatore JE, Aliev F, et al. Polygenic risk score prediction of alcohol dependence symptoms across population-based and clinically ascertained samples. Alcoholism: Clinical and Experimental Research. 2018;42(3):520–530.

10. Chang L-H, Couvy-Duchesne B, Liu M, et al. Association between polygenic risk for tobacco or alcohol consumption and liability to licit and illicit substance use in young Australian adults. Drug and alcohol dependence. 2019;197:271–279.

11. Jackson KM, Sartor CE. The natural course of substance use and dependence. In: Sher KJ, ed. The Oxford Handbook of Substance Use and Substance Use Disorders. New York, NY: Oxford University Press; 2016:67–134.

12. Malone SM, Taylor J, Marmorstein NR, McGue M, Iacono WG. Genetic and environmental influences on antisocial behavior and alcohol dependence from adolescence to early adulthood. Development and Psychopathology. 2004;16(4):943–966.

13. Bergen SE, Gardner CO, Kendler KS. Age-Related Changes in Heritability of Behavioral Phenotypes Over Adolescence and Young Adulthood: A Meta-Analysis. Twin Research and Human Genetics. 2007;10(3):423–433. doi:10.1375/twin.10.3.423

14. Vrieze SI, Hicks BM, Iacono WG, McGue M. Decline in genetic influence on the co-occurrence of alcohol, marijuana, and nicotine dependence symptoms from age 14 to 29. American Journal of Psychiatry. 2012;169(10):1073–1081.

15. Belsky DW, Moffitt TE, Baker TB, et al. Polygenic Risk and the Developmental Progression to Heavy, Persistent Smoking and Nicotine Dependence: Evidence From a 4-Decade Longitudinal Study. JAMA Psychiatry. 2013;70(5):534–542. doi:10.1001/jamapsychiatry.2013.736

16. Li JJ, Cho SB, Salvatore JE, et al. The impact of peer substance use and polygenic risk on trajectories of heavy episodic drinking across adolescence and emerging adulthood. Alcoholism: clinical and experimental research. 2017;41(1):65–75.

17. Ksinan AJ, Su J, Aliev F, et al. Unpacking genetic risk pathways for college student alcohol consumption: the mediating role of impulsivity. Alcoholism: Clinical and Experimental Research. 2019;43(10):2100–2110.

18. Su J, Kuo SI-C, Meyers JL, Guy MC, Dick DM. Examining interactions between genetic risk for alcohol problems, peer deviance, and interpersonal traumatic events on trajectories of alcohol use disorder symptoms among African American college students. Dev Psychopathol. 2018;30(5):1749–1761. doi:10.1017/S0954579418000962

19. Kranzler HR, Zhou H, Kember RL, et al. Genome-wide association study of alcohol consumption and use disorder in 274,424 individuals from multiple populations. Nature communications. 2019;10(1):1–11.

20. Sanchez-Roige S, Palmer AA, Fontanillas P, et al. Genome-wide association study meta-analysis of the Alcohol Use Disorders Identification Test (AUDIT) in two population-based cohorts. American Journal of Psychiatry. 2019;176(2):107–118.

21. Iacono WG, Carlson SR, Taylor J, Elkins IJ, McGue M. Behavioral disinhibition and the development of substance-use disorders: findings from the Minnesota Twin Family Study. Development and psychopathology. 1999;11(4):869–900.

22. Keyes MA, Malone SM, Elkins IJ, Legrand LN, McGue M, Iacono WG. The enrichment study of the Minnesota twin family study: increasing the yield of twin families at high risk for externalizing psychopathology. Twin Research and Human Genetics. 2009;12(5):489–501.

23. Wilson S, Haroian K, Iacono WG, et al. Minnesota Center for Twin and Family Research. Twin Research and Human Genetics. Published online 2019:1–7.

24. McGue M, Zhang Y, Miller MB, et al. A genome-wide association study of behavioral disinhibition. Behavior genetics. 2013;43(5):363–373.

25. Chang CC, Chow CC, Tellier LC, Vattikuti S, Purcell SM, Lee JJ. Second-generation PLINK: rising to the challenge of larger and richer datasets. Gigascience. 2015;4(1):s13742–015.

26. Das S, Forer L, Schönherr S, et al. Next-generation genotype imputation service and methods. Nature genetics. 2016;48(10):1284–1287.

27. Vilhjálmsson BJ, Yang J, Finucane HK, et al. Modeling linkage disequilibrium increases accuracy of polygenic risk scores. The american journal of human genetics. 2015;97(4):576–592.

28. Curran PJ, Howard AL, Bainter SA, Lane ST, McGinley JS. The separation of between-person and within-person components of individual change over time: A latent curve model with structured residuals. Journal of consulting and clinical psychology. 2014;82(5):879.

29. Berry D, Willoughby MT. On the practical interpretability of cross-lagged panel models: Rethinking a developmental workhorse. Child development. 2017;88(4):1186–1206.

30. Wu W, Selig JP, Little TD. Longitudinal data analysis. In: Little TD, editor. Oxford handbook of quantitative methods, Vol 2: Statistical Analysis. New York, NY: Oxford University Press; p. 387–410

31. Kwok O, West SG, Green SB. The impact of misspecifying the within-subject covariance structure in multiwave longitudinal multilevel models: A Monte Carlo study. Multivariate Behavioral Research. 2007;42(3):557–592.

32. Sivo S, Fan X, Witta L. The biasing effects of unmodeled ARMA time series processes on latent growth curve model estimates. Structural Equation Modeling. 2005;12(2):215–231.

33. Price AL, Patterson NJ, Plenge RM, Weinblatt ME, Shadick NA, Reich D. Principal components analysis corrects for stratification in genome-wide association studies. Nature genetics. 2006;38(8):904–909.

34. Muthén LK, Muthén BO. Mplus user’s guide. 8th ed. Los Angeles, CA

35. Allison, P. D. In Missing Data. Millsap RE, Maydeu-Olivares A. The SAGE Handbook of Quantitative Methods in Psychology. Sage Publications; 2009.

36. Falk CF. Are robust standard errors the best approach for interval estimation with nonnormal data in structural equation modeling? Structural Equation Modeling: A Multidisciplinary Journal. 2018;25(2):244–266.

37. West SG, Taylor AB, Wu W. Model fit and model selection in structural equation modeling. Handbook of structural equation modeling. 2012;1:209–231.

38. Zhou H, Sealock JM, Sanchez-Roige S, et al. Genome-wide meta-analysis of problematic alcohol use in 435,563 individuals yields insights into biology and relationships with other traits. Nature Neuroscience. Published online 2020:1–10.

39. Hicks BM, Foster KT, Iacono WG, McGue M. Genetic and environmental influences on the familial transmission of externalizing disorders in adoptive and twin offspring. JAMA psychiatry. 2013;70(10):1076–1083.

40. Kendler KS, Prescott CA, Myers J, Neale MC. The structure of genetic and environmental risk factors for common psychiatric and substance use disorders in men and women. Archives of general psychiatry. 2003;60(9):929–937.

41. Krueger RF, Hicks BM, Patrick CJ, Carlson SR, Iacono WG, McGue M. Etiologic connections among substance dependence, antisocial behavior, and personality: modeling the externalizing spectrum. Published online 2009.

42. Khera, A. V. et al. Genetic risk, adherence to a healthy lifestyle, and coronary disease. N. Engl. J. Med. 2016; 375, 2349–2358.

43. Lambert SA, Abraham G, Inouye M. Towards clinical utility of polygenic risk scores. Human molecular genetics. 2019;28(R2):R133–R142.

44. Torkamani A, Wineinger NE, Topol EJ. The personal and clinical utility of polygenic risk scores. Nature Reviews Genetics. 2018;19(9):581.

45. Driver, M. N., Kuo, S. I.-C., & Dick, D. M. Genetic feedback for psychiatric conditions: Where are we now and where are we going. American Journal of Medical Genetics Part B: Neuropsychiatric Genetics; 2020.

46. Mostafavi H, Harpak A, Agarwal I, Conley D, Pritchard JK, Przeworski M. Variable prediction accuracy of polygenic scores within an ancestry group. Elife. 2020;9:e48376.

47. Martin AR, Kanai M, Kamatani Y, Okada Y, Neale BM, Daly MJ. Clinical use of current polygenic risk scores may exacerbate health disparities. Nature genetics. 2019;51(4):584.

48. Rose RJ, Dick DM, Viken RJ, Kaprio J. Gene-environment interaction in patterns of adolescent drinking: regional residency moderates longitudinal influences on alcohol use. Alcoholism: Clinical and Experimental Research. 2001;25(5):637–643.

